# The duration and predictability of heatwaves shape host–parasite interactions under thermal stress

**DOI:** 10.64898/2026.02.27.708545

**Authors:** Viktoria Rozmann, Floriane O’Keeffe, Michael Officer, Pepijn Luijckx, Jeremy J. Piggott

**Affiliations:** Department of Zoology, School of Natural Sciences, Trinity College Dublin, Dublin, Ireland

**Keywords:** global change, host, disease, parasite, temperature variability, *Daphnia magna*, *Ordospora colligata*, *Hamiltosporidium tvaerminnensis*

## Abstract

Anthropogenic climate change is expected to increase not only mean temperatures but also the magnitude and pattern of thermal variability, including the frequency, duration, and predictability of extreme events. While the effects of elevated mean temperatures on disease dynamics are well studied, far less is known about how different patterns of temperature variability shape host–parasite interactions, despite clear theoretical predictions from the climate variability hypothesis. Here, we used a controlled experimental system (*Daphnia magna* and two microsporidian parasites, *Ordospora colligata* and *Hamiltosporidium tvaerminnensis*) to disentangle the effects of thermal variability structure from mean temperature. Across two experiments, we exposed hosts to cyclic (predictable) and random (unpredictable) heatwaves under both non-stressful and stressful mean temperature regimes. Contrary to predictions from the climate variability hypothesis, temperature variability did not uniformly increase infection risk. Instead, infection outcomes depended on the interaction between mean temperature, duration of the heatwaves, the pattern of thermal variation, and parasite identity. Under non-stressful mean temperatures, thermal variability had negligible effects on infection. In contrast, under stressful mean temperatures, parasites responded in distinct ways: *H. tvaerminnensis* infection was sensitive to the predictability of heat events, whereas *O. colligata* responded primarily to heatwave duration. These results demonstrate that climatic variability can differentially alter host–parasite interactions rather than exerting consistent directional effects on disease. By showing that parasite-specific sensitivities to the pattern of thermal variation emerge under thermal stress, our study highlights a mechanism by which increasing climatic variability may reshape parasite communities and disease outcomes in a warming world.

## Introduction

The COVID-19 pandemic and the occurrence of extreme weather events around the globe demonstrate the devastating impacts of both climate change and infectious diseases on food security (Ristaino et al., 2021), human health (Barrett et al., 2015; Newell & Dale, 2021) and ecosystem services (Mooney et al., 2009). While climate change and disease outbreaks can independently affect biodiversity (Garcia et al., 2014), lead to economic losses (Batten, 2018), and alter evolutionary trajectories (Parmesan, 2006), understanding their combined effects is crucial. Indeed, evidence is mounting that rising global temperatures are altering disease spread and causing more frequent outbreaks (Kirk et al., 2020; Lafferty, 2009; Rohr et al., 2011). For example, rising seawater temperatures are understood to be responsible for the increased prevalence of sea star wasting disease, which has led to the collapse of the sunflower star (*Pycnopodia helianthoides*) a keystone predator in marine ecosystems, leading to cascading effects that threaten biodiversity (Harvell et al., 2019). However, climate change is not only expected to raise mean temperatures but also to alter the variability and predictability of temperature regimes (Vasseur et al., 2014). These multifaceted changes can influence disease dynamics through their effects on both hosts and parasites, yet the underlying mechanisms and generality of these responses remain poorly understood (Claar & Wood, 2020). Disentangling how thermal variability affects host–parasite interactions is therefore essential for understanding disease outcomes under future climate scenarios.

Global change is anticipated to modify not only average temperatures but also induce shifts in temperature variability (Intergovernmental Panel on Climate Change (IPCC), 2023; Schär et al., 2004; Vasseur et al., 2014). Prior analyses underscore multiple unresolved questions pertaining to the occurrence of heatwaves, temperature fluctuations, and their impact on infectious diseases and parasites (Paaijmans et al., 2010; Raffel et al., 2013). The most urgent of these questions include whether sudden temperature spikes or fluctuations are distinct from gradual temperature increases (Rohr et al., 2013), and whether host–parasite interactions depend critically on the predictability of the temperature variation. Recent studies have demonstrated that the impact of heatwaves on diseases may be complex, and distinct from slowly rising mean temperatures (Kunze, Luijckx, et al., 2022; Lian et al., 2023). Indeed, heatwaves frequently yield diverse outcomes; for instance, in the system of the diamondback moth (*Plutella xylostella*) and its specialist parasitoid wasp (*Diadegma semiclausum*), a 10-degree temperature surge resulted in decreased parasitoid development, while a smaller 5-degree rise led to enhanced parasitoid growth (Schreven et al., 2017). This complexity arises not only due to the differential effects of temperature variation on various traits of hosts and parasites but also due to differences in the duration, intensity, and frequency of a heatwave which can determine whether disease prevalence is reduced or heightened (McCartan et al., 2025; Schreven et al., 2017). Various forms of temperature fluctuations, including daily temperature shifts, can similarly either amplify or diminish disease transmission (Kunze, Luijckx, et al., 2022; Paaijmans et al., 2010). A deeper understanding of the mechanisms underpinning the effects of thermal variation, and of whether the predictability of these fluctuations plays a critical role, are essential for advancing our understanding of host–pathogen interactions and disease epidemiology.

One proposed framework for understanding the complex impact of thermal variation is the Metabolic Theory of Ecology (MTE) (Brown et al., 2004), which links physiological processes such as growth, reproduction, and infection rates to body size and temperature-dependent metabolic rates. Building on this foundation, the climate variability hypothesis posits that increased, unpredictable temperature variability may temporarily favour parasites and pathogens over their hosts, owing to their smaller size and inherently faster metabolic rates (Raffel et al., 2013; Rohr et al., 2013), allowing them to acclimate more rapidly to environmental fluctuations than their hosts. Recent empirical studies provide support for this hypothesis; for instance, Kunze, Luijckx, et al. (2022), and Krichel et al. (2023) observed increased infection success in *Daphnia magna* and its parasite *Ordospora colligata* under fluctuating temperature. Accordingly, host–parasite systems with size-based metabolic differences, such as *D. magna* and its microparasites, offer a powerful model for experimentally testing how temperature variability shapes disease outcomes, and whether parasites consistently gain an advantage under fluctuating thermal conditions.

Here we examine the effect of predictable and unpredictable temperature fluctuations in two experiments using *Daphnia magna* and two of its microsporidian parasites, *Ordospora colligata* and *Hamiltosporidium tvaerminnensis*. Both microcosm experiments were designed to test how different patterns of temperature variation, including several predictable (cyclic) and unpredictable (random) fluctuations, affect host and parasite performance, with one conducted within a non-stressful and the other under a stressful temperature range. Our experiments specifically test how different forms of temperature variation influence disease outcomes, the potential advantage of unpredictable fluctuating temperatures for parasites, the consistency of responses across both parasites, and under what conditions thermal stress leads to reduced parasite performance. We hypothesise under stressful temperature conditions the parasite’s advantage may diminish because fluctuations that exceed the parasite’s thermal tolerance can impair key physiological processes. This expectation is based on the thermal stress hypothesis, which posits that parasites with narrower thermal performance ranges than their hosts are more susceptible to performance declines under such conditions (Paull et al., 2015).

## Materials and methods

### Study system

The two experiments used the same study system, *Daphnia magna* and two of its microsporidian parasites, *Ordospora colligata* and *Hamiltosporidium tvaerminnensis*. *D. magna* naturally encounters spores of both parasites across its range. These spores are ingested by the host *D. magna* during filter feeding (Ebert, 2005), after which they germinate in the gut epithelium, and discharge their polar filament (a tightly coiled tube) which transfers the nucleus and sporoplasm to the host (Han et al., 2022). While *O. colligata* infects and grows intracellularly within cells of the gut, forming clusters of up to 64 spores, *H. tvaerminnensis* exclusively infects the fat and ovarian cells (Haag et al., 2011). The shedding mechanisms for these two parasites also differ. *O. colligata* continuously releases its spores into the environment through the host’s faeces, facilitating horizontal transmission (Ebert et al., 2000). In contrast, *H. tvaerminnensis* exhibits both vertical and horizontal transmission, upon the death of the host. The genotypes of the host and the parasites used in this experiment were kept in the lab for over 5 years but were originally collected from the Tvärminne Archipelago in Finland (*D. magna* genotype Fi-Oer-3-3, *O. colligata* isolate OC3 and *H. tvaerminnensis*).

### Experiments

The first experiment, testing the temperature variability hypothesis in a non-stressful temperature range (16–22°C) for both *O. colligata* and *H. tvaerminnensis*, lasted 126 days. We compared five predictable (cyclic) and three unpredictable (random) temperature treatments for both parasites and a placebo control treatment, for a total of 24 treatments. The second experiment followed a similar design to the first but was conducted under a stressful temperature range (20–26°C) and lasted 78 days. It included three predictable and three unpredictable fluctuating temperature treatments for both the parasites and placebo controls (18 treatments in total). Additional constant temperature controls were also included in both experiments.

### Experiment preparation

The same procedures were followed in preparation for both experiments: ∼ 500 adult *D. magna* of genotype Fi-Oer-3-3 were grown for four weeks under standardised conditions to minimise maternal effects. Groups of 10-12 female animals per 400 mL microcosm with 200 mL of artificial Daphnia medium ADaM (Ebert, 2005) were kept at 20°C, transferred to a new microcosm with fresh medium twice a week, and fed *ad libitum* with batch cultured *Scenedesmus sp.* algae grown in WC medium (Kilham et al., 1998). The animals were transferred to new microcosms four days before the start of the experiment to ensure the absence of juveniles. Any juveniles born within the subsequent four days (96 hours) were gathered, sexed using a dissecting microscope (8 to 12x magnification), and females were individually placed in 100 mL microcosms containing 60 mL of ADaM while males were discarded. Microcosms with juvenile females were arranged into trays, placed in the designated water baths, and rotated daily to minimize positional effects. For Experiment 1 we used a uniform block design with one replicate per fluctuating temperature in each tray. However, for Experiment 2 an alternative design was chosen, with 8 replicates of *O. colligata* and *H. tvaerminnensis* of one fluctuating temperature treatment per tray and three trays per temperature treatment, as the uniform block design greatly increased handling time per sample and led to time constraints. Spores for both parasites were propagated by allowing infections to spread through populations of *D. magna* maintained in either 5 L buckets for *O. colligata* or 2 L glass beakers for *H. tvaerminnensis*. These populations were fed *Scenedesmus* sp. algae three times per week, with approximately 1,500 million algae provided for *O. colligata* populations and 750 million for *H. tvaerminnensis* populations.

#### Parasite spore preparation

Spore doses were selected based on established protocols and pilot studies for each host-parasite combination (see Kunze et al. (2022) and Kirk et al. (2018) for *O. colligata*; and Visozo, Lass and Ebert (2005) and Krebs, Routtu and Ebert (2017) for *H. tvaerminnensis*). For *H. tvaerminnensis*, spores were prepared by homogenizing individuals from the infected population using a mortar and pestle, and the resulting spore solution was quantified with a Neubauer improved haemocytometer under phase-contrast microscopy. For *O. colligata*, infected individuals from a stock population with known infection prevalence and spore burden (confirmed via phase-contrast microscopy on a subsample) were similarly homogenized. Spore solutions for both parasites were then diluted to a concentration of 50,000 spores per mL. A placebo dose was prepared by homogenising uninfected individuals from the same *D. magna* strain used in the experiment (Fi-Oer-3-3), following the same procedure as for the spore doses. On the day of exposure (defined as day 0), each individual received 1 mL of the corresponding spore suspension or placebo based on their assigned treatment. Animals remained in the exposure environment for 7 days before being transferred to fresh ADaM.

### Temperature manipulation

Temperature of the microcosms was manipulated using water baths interfaced with a temperature controller (Inkbird ITC-308D or ITC-310T-B) which was connected to aquarium chillers (Hailea HC150A, DC300 or DC750) and heating rods (EHEIM JÄGER 300W). Microjet pumps (NewaMicro MC450 or Oase OptiMax 500) were used to circulate the water and maintain an even temperature. Temperature fluctuations were implemented by moving individual jars between trays (Experiment 1), or whole trays, containing 24 microcosms (Experiment 2), to a water bath of the correct temperature. In total we set up 6 water baths at 16°C and 6 at 22°C for Experiment 1, and the same number of water baths at 20°C and at 26°C for Experiment 2, with additional constant temperature controls at the mean temperature (at 17.4°C for Experiment 1 and 21.5°C for Experiment 2).

### Experimental design

#### Experiment 1

Microcosms containing individual juvenile females of *D. magna* were exposed to either spores of *O. colligata, H. tvaerminnensis*, or a placebo comprised of crushed uninfected *D. magna* individuals from the Fi-Oer-3-3 clonal line. After a 10-day acclimation period at 16°C, the animals were exposed to either cyclic (predictable) or random (unpredictable) temperature shifts between 16°C and 22°C. The cyclic temperature shifts varied in duration, lasting 10, 5, 2.5, 1.25, and 1 day, and were repeated 2, 4, 8, 16, and 20 times, respectively, over an 87-day span (Figure 1, left column). The random temperature fluctuation had a duration of 2.5 days each, occurring in a random pattern that ensured a minimum 2.5-day recovery period between the fluctuations. Each parasite-temperature treatment combination was applied to 24 replicates, for a total of 576 individuals. All temperature treatments were designed to keep the mean temperature at 17.4°C, over the 87-day period. After this period, the 87-day cycle was restarted with new seeds for the randomly fluctuating temperatures. A parallel experiment (O’Keeffe, 2024) using identical protocols, which was designed to estimate thermal performance of both *O. colligata* and *H. tvaerminnensis* provided responses of both parasites at five constant temperatures including the mean temperature of 17.4°C (in 24 replicates).

**Figure 1:**
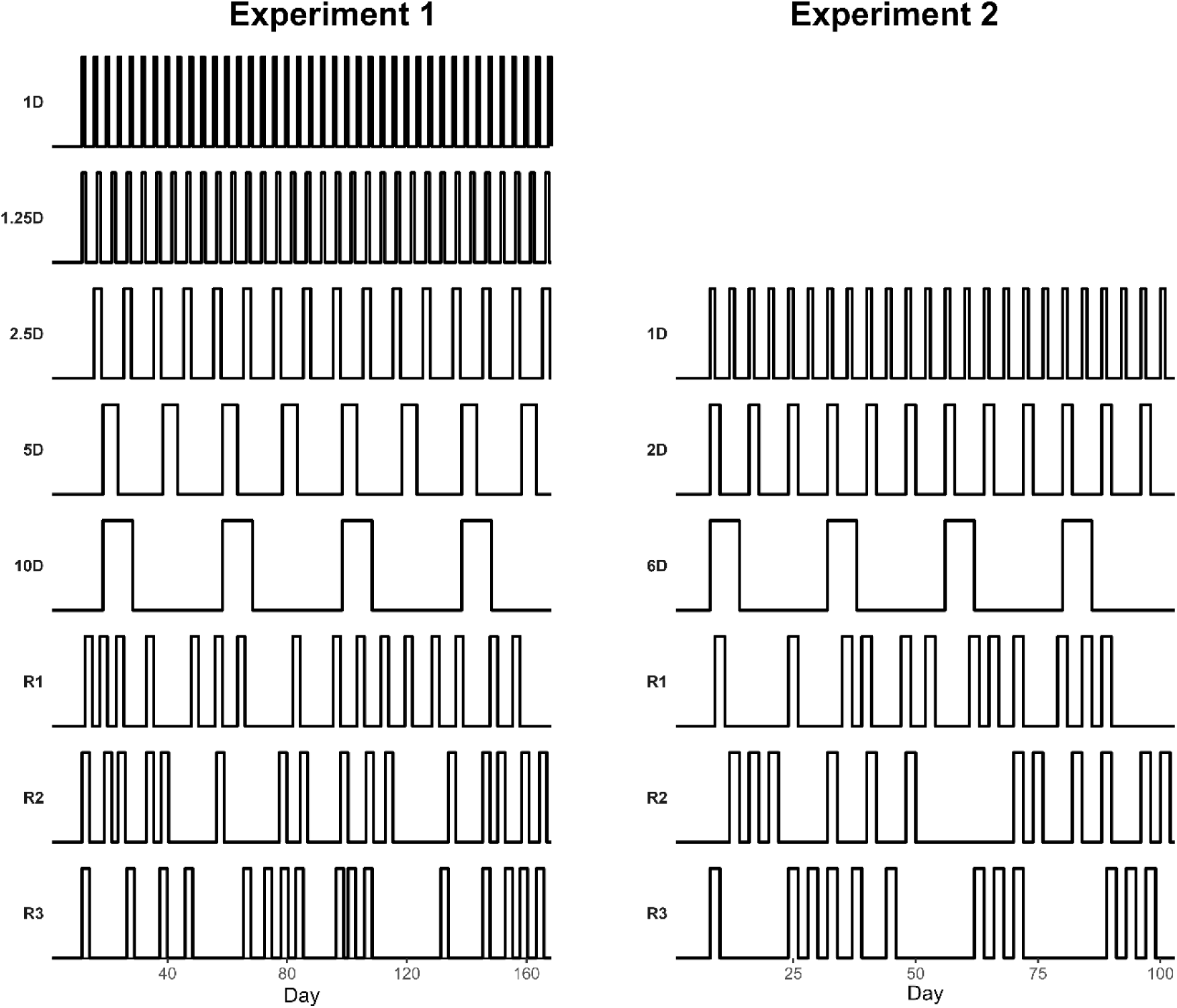
Temperature treatments of Experiment 1 (left; non-stressful conditions) and Experiment 2 (right; stressful conditions). Experiment 1 had five predictable fluctuating temperatures, and three unpredictable ones. Experiment 2 had three predictable and three unpredictable treatments. A simultaneous experiment measured thermal performance across five temperatures during Experiment 1, and these values are used as average controls while Experiment 2 included controls at the mean, minimum and maximum temperature.

#### Experiment 2

Experiment 2 had a similar design to Experiment 1, however the mean temperature used was higher (21.5°C). For this experiment we used three trays with 24 microcosms with individual *Daphnia* for each of the six temperature treatments instead the uniform block design in Experiment 1. Each microcosm was exposed to either parasite or placebo and subjected to either cyclic (predictable) or random (unpredictable) temperature fluctuations after a 7-day acclimation period at 20°C. Temperature shifts occurred between 20°C and 26°C. Three cyclic treatments exposed individuals to higher temperatures for 1, 2, or 6 days, occurring 12, 6, and 2 times, respectively, over a 55-day period, while unpredictable random temperature fluctuation treatments lasted 2 days each, with a total of six shifts over the same period. Similar to Experiment 1, there were 24 replicates in each parasite-temperature treatment combination and constant temperature controls were also included, for a total experimental size of 648 individuals.

### Experiment maintenance

Animals in both experiments were kept under a 16:8 light–dark photoperiod and were transferred to new microcosms with fresh artificial Daphnia medium ADaM (Ebert et al., 1998) twice a week (the first transfer occurring 7 days post-infection) to prevent the accumulation of waste and offspring produced during the experiment. Each animal was fed with 1 mL *Scenedesmus* algae 4 times a week, starting with a density of 5 million cells/mL, which gradually increased to 12 million cells/mL in week 1, remaining constant thereafter. Experiment 1 ran for 126 days, the last 14 surviving animals were terminated on day 126 while experiment 2 lasted 78 days, with the last 9 animals terminated on day 78.

### Measurements

*D. magna* were checked daily for mortality and deceased individuals were subsequently examined for infections. Dead animals were dissected on a microscope slide in 100 µL of water under a stereo microscope. Infection rate was defined as the proportion of examined individuals in which a parasite was detected. *O. colligata* infection and spore burden, defined as the number of spore clusters present in the gut of an infected animal were determined using phase-contrast microscopy at 400× magnification. The presence of *H. tvaerminnensis* was assessed by crushing and homogenising the dissected animal on the slide, and spore burden was quantified as the number of spores per individual by analysing 12.5 µL of the resulting liquid on a haemocytometer under phase-contrast microscopy at 400x magnification.

### Statistical analyses

All statistical analyses were performed using R (version 4.4.1) and RStudio (build 764, “Chocolate Cosmos”). All analyses were performed using a generalised linear model (GLM) for Experiment 1, and GLMs and generalised linear mixed models (GLMMs) for Experiment 2 with appropriate error distributions checked for overdispersion where relevant. Variable significance was assessed by model comparison using an analysis of deviance for GLMs and Type II Wald Chi squared tests for GLMMs. Animals that died before the 8^th^ day of the experiment were excluded from analysis since determination of infection status is not reliable at this stage (n = 14 in Experiment 1 and n = 28 in Experiment 2). For Experiment 1 parasite fitness measures were analysed using a negative binomial distribution for spore burden and binomial distribution for infection rates, separately for the two parasites. Host longevity was assessed using a GLM with a negative binomial distribution with temperature treatments and parasite as explanatory variables (days since exposure ∼ treatment*parasite). For Experiment 2 we tested if adding “tray” to the model (GLMM) as a random effect improves the model fit and used GLMs wherever the random effect was estimated to be negligible. Model formulations and used distributions were similar to Experiment 1 (that is, negative binomial for host longevity and spore burden and binomial for infection rates). For host longevity, the final model was (days since exposure ∼ treatment + (1|tray2)) while models for infection and burden were formulated as (infection status ∼ treatment) and (number of spores ∼ treatment), respectively. For both Experiment 1 and 2 we estimated marginal means (EMMs) (emmeans package, version 1.10.4) when models detected significant effects.

## Results

The differing mean temperatures of the two experiments revealed distinct effects on host-parasite dynamics, with differences observed only under the higher, stressful mean temperature in Experiment 2. Under the non-stressful mean temperature (17.4 °C) of Experiment 1, animals infected with *H. tvaerminnensis* died earlier than *O. colligata* infected or control individuals (GLM, df = 2, p<0.001). However, parasite burden and infection rates did not differ among treatments, and fluctuating temperature regimes had no detectable effects on host fitness, regardless of cycle duration or predictability (GLM, all p > 0.05). The higher mean temperature (21.5°C) of Experiment 2 was stressful to the host and both of the parasites, resulting in reduced fitness (mean host longevity: 86 days in Experiment 1 vs 42 days in Experiment 2; mean *O. colligata* spore clusters: 494 vs 37.5; mean *H. tvaerminnensis* spore load: 20.5 vs 16.8 million). Moreover, in contrast to Experiment 1, Experiment 2 showed that under stressful conditions, both the specific temperature pattern and the duration of the cycle affected the outcome; see Figure 2 and Table 1 for all results.

**Figure 2:**
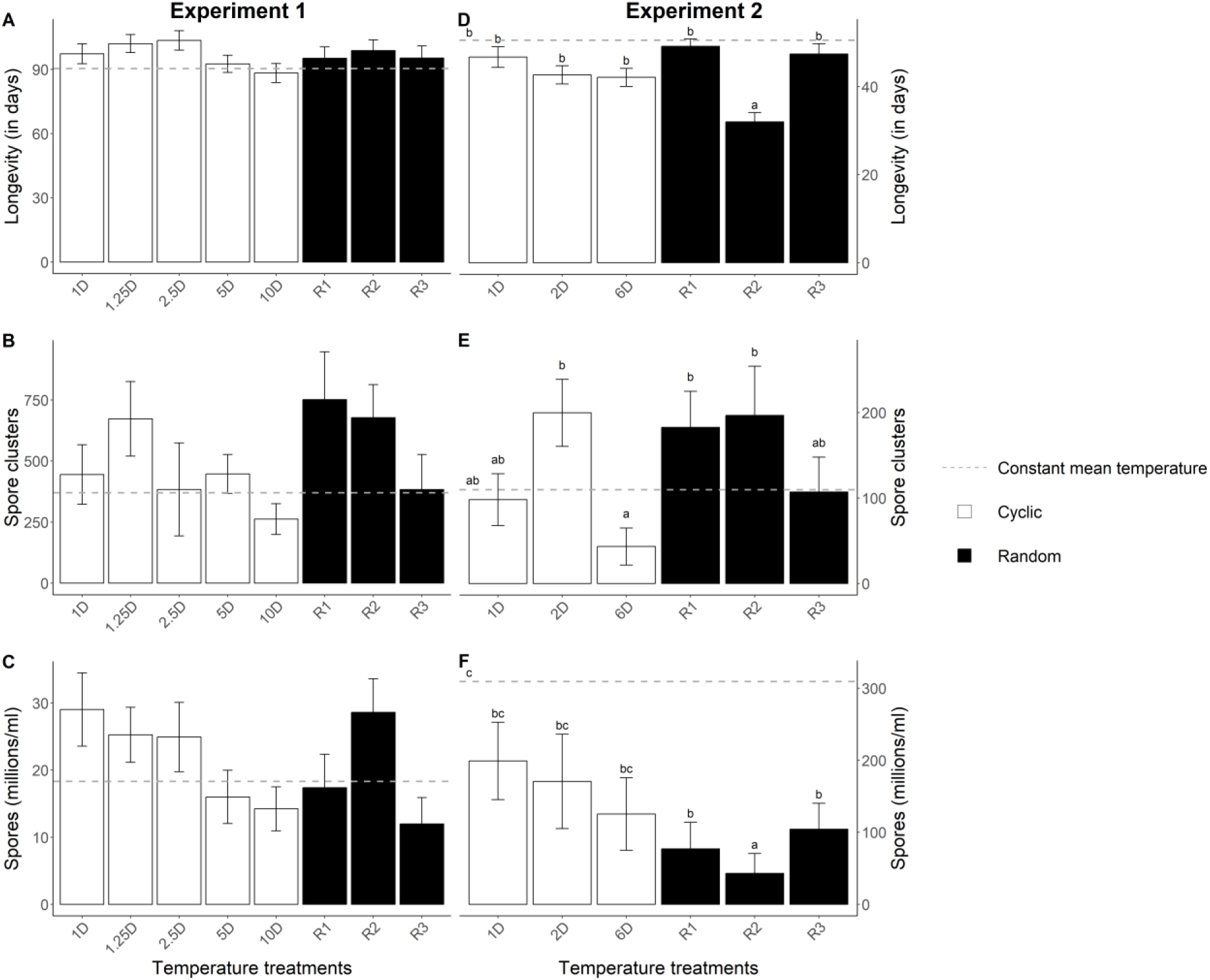
Longevity of the unexposed control animals in days (A) and longevity of all animals (D), average of O. colligata spore clusters and average of H. tvaerminnensis spores by temperature treatment in the two experiments (Experiment 1 in the left column and Experiment 2 in the right column). Constant mean temperatures are denoted by dotted lines; cyclic temperatures are represented in white and random temperatures are represented in black. Error bars represent standard error. Different letters above bars indicate statistically distinct groups based on pairwise multiple comparisons (see Tables S1, S2, and S3). Treatments sharing a letter are not significantly different from one another.

**Table 1:** Summaries of generalised linear models for Experiment 1 and 2.

| Experiment 1 | factor | Df | deviance | Resid. Df | Resid. Dev | p |
| --- | --- | --- | --- | --- | --- | --- |
| Host longevity | treatment | 8 | 10.355 | 604 | 696.94 | 0.241 |
|  | parasite | 2 | 46.808 | 602 | 650.14 | 6.85E-11 *** |
|  | treatment x parasite | 16 | 7.857 | 586 | 642.28 | 0.953 |
| *O. colligata* infection | treatment | 8 | 5.6667 | 193 | 153.22 | 0.6845 |
| *O. colligata* burden | treatment | 8 | 13.912 | 120 | 149.41 | 0.08408 . |
| *H. tvaerminnensis* infection | treatment | 8 | 7.5055 | 200 | 280.12 | 0.4832 |
| *H. tvaerminnensis* burden | treatment | 8 | 10.218 | 103 | 127.54 | 0.2501 |

| Experiment 2 | factor | Df | Chisq/<br>deviance | Resid. Df | Resid. Dev | P |
| --- | --- | --- | --- | --- | --- | --- |
| Host longevity* | treatment | 8 | 107.2078 | - | - | <2e-16 *** |
|  | parasite | 2 | 3.2393 | - | - | 0.198 |
|  | treatment x parasite | 16 | 15.5851 | - | - | 0.4823 |
| *O. colligata* infection | treatment | 8 | 22.862 | 183 | 203.38 | 0.003546 ** |
| *O. colligata* burden | treatment | 8 | 52.611 | 121 | 152.65 | 1.28E-08 *** |
| *H. tvaerminnensis* infection | treatment | 8 | 23.007 | 197 | 253.1 | 0.003355 ** |
| *H. tvaerminnensis* burden | treatment | 8 | 50.721 | 112 | 157.19 | 2.97E-08 *** |

\*Type II Wald Chi squared tests

In Experiment 2, host and parasite fitness differed between treatments, although both parasites showed different outcomes. Host mortality varied by treatment (GLM, df = 8, p<0.001) and was particularly high in one of the unpredictable treatments (R2), which differed from the constant temperature control as well as all other predictable and unpredictable fluctuations (all pairwise contrasts for R2 p<0.0168, see Table S1). This treatment experienced three cycles of elevated temperature in rapid succession during the first 23 days of the experiment, with a large decrease in survival beginning after the second cycle and continuing until a few days after the third cycle (Figure 3). Other unpredictable treatments experienced only a single event of elevated temperature, with a long recovery period over this time.

**Figure 3:**
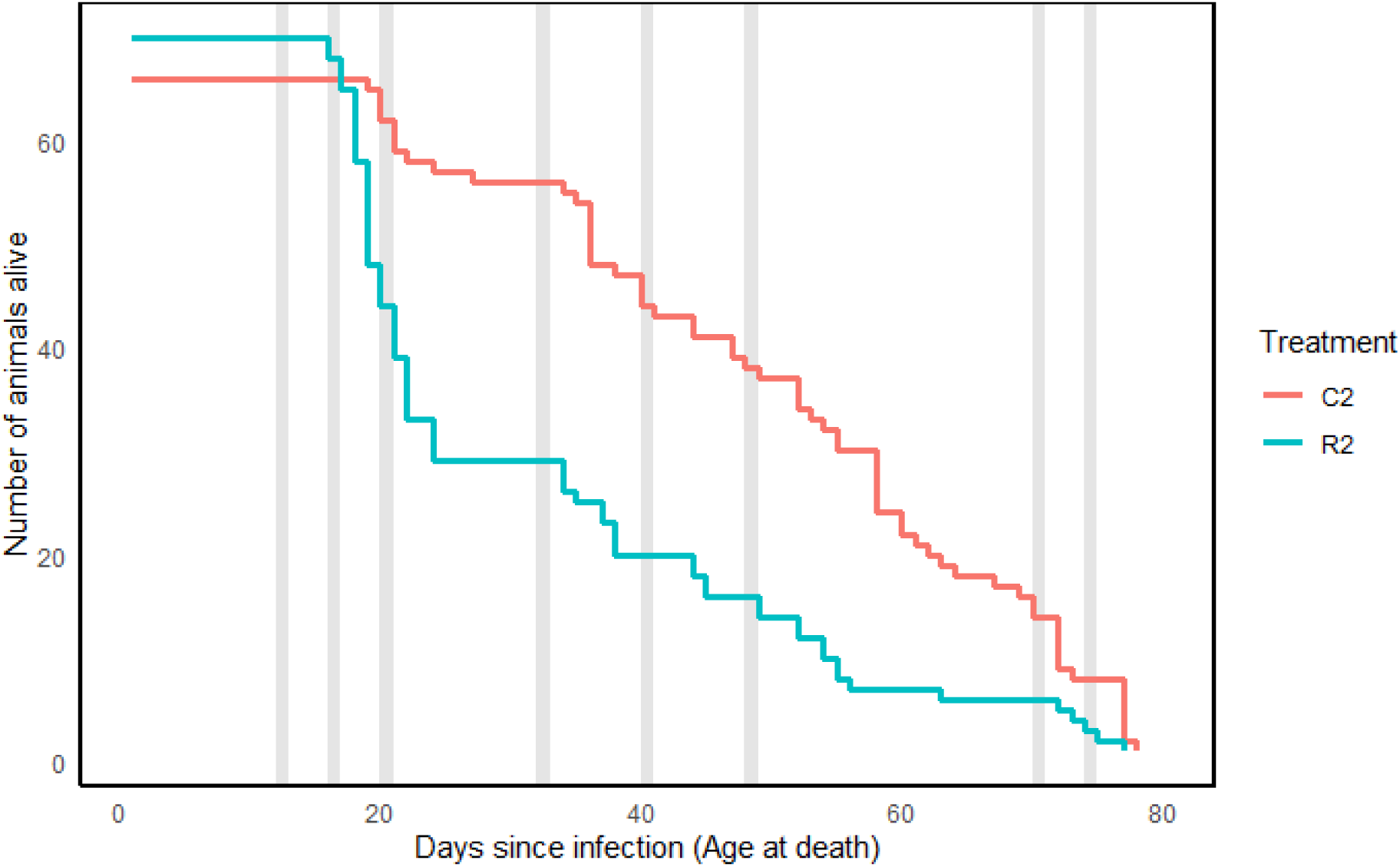
Mortality in the R2 unpredictable temperature treatment (blue) in Experiment 2, compared to the mean temperature control (red, C2). Gray bars indicate the timing of the heat events.

Infection rates were affected by temperature treatments (GLM, df = 8, p<0.01 for both parasites). For *O. colligata*, only the high constant temperature had a significant effect on infection rates (p<0.05, see Supplemental Figure 1). For *H. tvaerminnensis*, the overall effect of treatments was significant, but pairwise comparisons only showed trends toward differences (p = 0.051), with treatment R2 tending to differ from the constant temperature control (Supplemental Figure 2).

The spore production of *H. tvaerminnensis* was generally lower in fluctuating environments, with all treatments lower than the constant average temperature control (although not significantly for all treatments), see Table S2. This was especially true for two of the unpredictable treatments, R1 and R2, which differed from the average temperature control, respectively (GLM, df = 8, both p<0.01). Among the three unpredictable treatments, R2 also showed a marginal difference from the predictable treatment with the same cycle length (R2 vs D2, p=0.08).

Although unpredictable temperature variation may have led to lower spore burden for *H. tvaerminnensis*, *O. colligata* was primarily influenced by the duration of the temperature events. The number of spore clusters that grew within the host was lower when duration of the elevated temperature was 6 days. Indeed, this treatment was marginally different from the constant temperature controls and the heat events that lasted 2 days (both predictable and unpredictable), see Table S3. Unlike *H. tvaerminnensis*, *O. colligata* didn’t seem to be universally stressed by fluctuating temperatures with all other treatments showing either the same or slightly higher parasite burden than the average temperature control. Infection rates were similar across all treatments (except the high constant temperature), supporting that for *O. colligata* in the treatments used in our experiment were generally not stressed by fluctuating temperatures except when these were 6 days long.

## Discussion

Our results demonstrate that the effects of temperature fluctuations on host–parasite interactions depend strongly on the thermal context. Under non-stressful temperature conditions, temperature fluctuations did not have any effect on parasite burden or host mortality rates. However, both host and parasite performance declined under stressful temperatures, showing an effect of thermal stress on the model system. Notably, only some patterns of thermal variation affected the three species studied: *O. colligata* was sensitive to the duration of the heat event, while both the host and *H. tvaerminnensis* lost fitness for certain unpredictable temperature patterns. These results highlight that the effects of temperature variation are context-dependent and species specific, with important implications for understanding parasite dynamics under climate-induced temperature fluctuations. Because thermal variability under non-stressful conditions had no detectable effects, we focus the remainder of the discussion on Experiment 2, which examined host–parasite dynamics under stressful temperature regimes.

Thermal fluctuations had the greatest impact when temperatures approached or exceeded the thermal limits of the organisms, leading to reduced fitness across several fluctuating treatments for both the parasites and the *Daphnia* host. Studies on the *O. colligata*–*Daphnia* system support these findings, showing that both fluctuating temperatures and heatwave events reduce fitness at thermal extremes due to stress associated with high temperatures (Kunze, Luijckx, et al., 2022; McCartan et al., 2025). In contrast, previous work using single heatwaves or diurnally fluctuating temperatures suggested that *H. tvaerminnensis* did not suffer reductions in fitness due to thermal stress, potentially due to its broader thermal tolerance than the host (O’Keeffe, 2024); here, we show that persistent, long-term fluctuations are clearly stressful in line with the thermal stress hypothesis (Paull et al., 2015). Indeed, that temperature fluctuations can alter host-parasite interactions has been recorded in numerous host-parasite systems (see Claar & Wood, 2020 for a review). For example, temperature fluctuation near the thermal maximum have been shown to reduce malaria fitness (Paaijmans et al., 2010), however, thermal stress can also reduce host immune function and provide an advantage to the pathogen (Seppälä & Jokela, 2010). While the thermal stress hypothesis thus seems to apply to the *Daphnia*-parasite system, no support was found for the climate variability hypothesis.

Contrary to the expectations of the climate variability hypothesis (Raffel et al., 2013; Rohr et al., 2013), we found no increase in parasite performance under fluctuating temperatures. Under this hypothesis, parasites are expected to outperform their hosts during unpredictable temperature changes because their smaller size leads to faster metabolic rates, enabling them to acclimate more quickly than their larger hosts (Raffel et al., 2013). Although previous studies using *O. colligata* have invoked this theory to explain a four-fold increase in parasite burden following a heatwave (Kunze, Luijckx, et al., 2022) and elevated endemic prevalence after short-term fluctuations (Krichel et al., 2023), our experiment—which explicitly tested both predictable and unpredictable thermal variation—did not support this hypothesis. Notably, all experiments, including those of Krichel et al., (2023) and Kunze, Luijckx, et al., (2022), were conducted using the same *D. magna* clone originating from the Tvärminne Archipelago in Finland. In this habitat, *D. magna* inhabits rockpools in which temperatures can fluctuate (Legrand et al., 2018). As a result, the animals may exhibit high phenotypic plasticity in response to changing environmental conditions, making them resilient to environmental variability itself and potentially obscuring support for the climate variability hypothesis in our experiment, although additional mechanisms may still be required to explain the increases in parasite load observed in both previous experiments. In addition, empirical tests of the climate variability hypothesis are inherently challenging, as temperature fluctuations simultaneously impose direct thermal stress on both host and parasite and affect multiple physiological processes (Altman et al., 2016; Paull et al., 2015; Scharsack et al., 2021). This makes it difficult to disentangle effects driven specifically by differential acclimation rates from those arising from thermal stress per se or other temperature-dependent mechanisms. However, support for the hypothesis exists in other systems, for example unpredictable temperature fluctuations decreased Cuban treefrog (*Osteopilus septentrionalis*) resistance to *Batrachochytrium dendrobatidis*, a pathogenic chytrid fungus (Raffel et al., 2013). While our work does not support the climate variability hypothesis, it does show that not all thermal variation leads to the same outcomes and that predictability and pattern of the temperature variation can matter.

Some of the unpredictable fluctuating-temperature regimes reduced fitness for the host (R2) and for *H. tvaerminnensis* (R2, R3) when temperatures fluctuated near the thermal maximum (Experiment 2). One of the unpredictable fluctuating-temperature regimes (R2) had heightened mortality during three heat events that occurred within 10 days early in the experiment, potentially providing limited time for acclimation (Peck et al., 2014) or recovery (Müller et al., 2018). Species can require several days to adjust to new thermal conditions (Weldon et al., 2011) and in *Daphnia*, acclimation can take up to a week, with recovery periods of 2–4 days following heat stress (Müller et al., 2018) before performance returns to normal. The reduced performance observed in this unpredictable temperature regime may therefore reflect insufficient time for acclimatisation and recovery. Comparable patterns have been found in the tobacco hornworm (*Manduca sexta*), where multiple heatwaves, but not a single event, reduced pupal mass and where heatwaves occurring early or mid-development decreased body mass (Kingsolver et al., 2021). Similarly, in our system the occurrence of multiple heatwaves early in life could have led to a differential outcome as thermal fluctuations can have age-dependent effects (Salachan & Sørensen, 2017; Schou & Cornwallis, 2024). While other unpredictable treatments also experienced multiple heat events in a short period later in the experiment, their prior exposure to a heat event with sufficient time for recovery, hardening, or acclimatisation may have allowed them to cope better with future temperature events (Le Lann et al., 2021). Plastic responses to thermal fluctuation have been found for hosts, parasitoids (Le Lann et al., 2021), and parasites (Lee et al., 2016). For instance, both the host, the spotted stem borer *Chilo partellus*, and its endoparasitoid *Cotesia flavipes* showed increased survival in a heat event when pretreated with a short exposure to elevated temperature (Mutamiswa et al., 2018). While rapid fluctuations in temperature early in the experiment thus increased *Daphnia* mortality, they likely also indirectly influenced the performance of the *H. tvaerminnensis* parasite, lowering infection rates as well as spore burden.

Spore production of *H. tvaerminnensis* was reduced when temperatures fluctuated rapidly early in the experiment (treatment R2), likely in part because reduced host survival left the parasite with less time to grow. However, given that one of the other unpredictable treatments (R1) was also lower than average controls, this suggests that additional factors may also contribute to reduced performance in fluctuating environments. Indeed, unpredictable thermal environments may cause disturbances in the regulation of heat shock protein (HSP) expression (Angilletta Jr., 2009), and the cost of adopting the wrong strategy of HSP production can lower the fitness of the parasite. In addition, HSPs may play an important role on the host side, where unpredictable temperature fluctuations can induce elevated HSP expression. This pattern has been observed in qingbo fish (*Spinibarbus sinensis*) exposed to predictable, unpredictable and constant temperature regimes, where fish subjected to unpredictable temperature fluctuations exhibited the highest levels of *hsp70* expression (Fu et al., 2024). Another possible factor is, since parasite development is tightly synchronised with host cellular processes (Motta et al., 2023), that under random temperature regimes, thermal unpredictability may disrupt host circadian rhythms and immune responses, as has been observed in the red swamp crayfish (*Procambarus clarkii*) (Dong et al., 2015), creating unfavourable intracellular conditions for parasite development. However, why these effects were observed in *H. tvaerminnensis* but not *O. colligata*, remains unclear.

At higher mean temperatures, slower fluctuating regimes with long periods of unfavourable temperatures negatively affected the spore production of *O. colligata*. The 6-day heat event produced fewer spore clusters than the 2-day heat event; Raffel et al., (2013) predicted this behaviour, assuming that longer time periods between temperature shifts will result in lower pathogen growth rates. Also, longer exposure to stressful temperatures can have a larger impact than short term fluctuations, as seen in other study systems, including *Emiliania huxleyi* (Wang et al., 2019)*, Mytilus edulis* (Pansch & Hiebenthal, 2019) and phytoplankton communities (Kunze, Gerhard, et al., 2022). While exposure to high temperatures may induce the production of heat-shock proteins (Angilletta Jr., 2009), which can increase survival during subsequent high-temperature events, longer exposure (and consequently higher levels of heat-shock proteins) may reduce fitness (Kingsolver et al., 2015; Krebs & Feder, 1997). Indeed, overexpression of heat shock proteins leads to increased energetic costs and lower fitness in *Drosophila melanogaster* (Hoekstra & Montooth, 2013; Krebs & Feder, 1997). Another explanation for the difference between the long duration and shorter duration heat events can be the that host and parasite fitness components can respond differently to temperature across timescales. In particular, short-term increases in heat may temporarily enhance parasite performance, while prolonged heat exposure may push parasites beyond their thermal optima, reducing replication, survival, or overall infection intensity (McCartan et al., 2025; Ragonese et al., 2024). At the same time, longer heatwaves may allow hosts to acclimate physiologically or reallocate resources, further contributing to reduced parasite loads, as seen in the *D. magna* – *Pasteuria ramosa* system (Hector et al., 2024). With climate reddening, weather conditions are expected to shift more slowly and become more persistent, leading to longer and more sustained periods of high temperatures (Di Cecco & Gouhier, 2018). As our results demonstrate, prolonged heat exposure may impose additional stress on organisms even when mean temperatures remain stable underscoring the importance of temperature variability in shaping species interactions, particularly when hosts and parasites are affected differently.

### Conclusion/Implications

In this study we empirically tested the climate variability hypothesis in a host–parasite system exposed to cyclic and unpredictable temperature fluctuations. Although our results did not support the hypothesis, they reveal that the effects of temperature variability are contingent on parasite identity and the temporal characteristics of thermal stress. Under benign conditions temperature fluctuations had limited impact and while at higher, stressful temperatures, some patterns of temperature variation were similar for *D. magna* and its two parasites, others differed. Indeed, *Hamiltosporidium tvaerminnensis* infections depended on the pattern of temperature variation, whereas *Ordospora colligata* was primarily affected by the duration of heat events. Consistent with growing evidence that co-occurring pathogens can differ in their thermal sensitivity, these findings indicate that temperature variability may reshape parasite communities rather than uniformly increasing or decreasing disease risk. Moreover, altered thermal regimes can elevate host mortality and modify host–parasite interactions through multiple, context-dependent physiological and behavioural pathways, with consequences for virulence and transmission dynamics. Together, our results highlight that increasing climatic variability is unlikely to have uniform effects on disease outcomes, underscoring the challenge of predicting ecological responses to global change and the need to account explicitly for species interactions and the structure of thermal variation.

## Supporting information

All supplemental tables and figures

## Acknowledgements

We thank Dieter Ebert and Jürgen Hottinger for providing *Daphnia* used in this study.

