## Supplementary material for "The duration and predictability of heatwaves shape host–parasite interactions under thermal stress": All supplemental tables and figures

Supplements for Heatwaves reshape host–parasite interactions

*Table 1: Pairwise contrasts of estimated marginal means (log scale) from a negative binomial GLMM of overall longevity in Experiment 2.*
*P-values adjusted using the Benjamini–Hochberg method; 95% CI computed on log scale.*

| **Contrast** | **Log difference (LRR)** | **95% CI** | **adj. p** |
|---|---|---|---|
| 1D – 2D | 0.081 | [−0.12, 0.28] | 0.521 |
| 1D – 6D | 0.096 | [−0.11, 0.30] | 0.456 |
| 1D – C1 | −0.195 | [−0.40, 0.01] | 0.119 |
| 1D – C2 | −0.088 | [−0.29, 0.12] | 0.496 |
| **1D – C3** | **0.744** | **[0.53, 0.96]** | **<0.0001** |
| 1D – R1 | −0.060 | [−0.27, 0.15] | 0.640 |
| **1D – R2** | **0.374** | **[0.17, 0.58]** | **0.0010** |
| 1D – R3 | −0.018 | [−0.22, 0.19] | 0.885 |
| 2D – 6D | 0.015 | [−0.19, 0.22] | 0.885 |
| **2D – C1** | **−0.276** | **[−0.48, −0.07]** | **0.017** |
| 2D – C2 | −0.169 | [−0.37, 0.03] | 0.181 |
| **2D – C3** | **0.663** | **[0.45, 0.88]** | **<0.0001** |
| 2D – R1 | −0.141 | [−0.35, 0.07] | 0.281 |
| **2D – R2** | **0.293** | **[0.09, 0.50]** | **0.012** |
| 2D – R3 | −0.099 | [−0.30, 0.11] | 0.454 |
| **6D – C1** | **−0.292** | **[−0.50, −0.09]** | **0.012** |
| 6D – C2 | −0.184 | [−0.39, 0.03] | 0.149 |
| **6D – C3** | **0.648** | **[0.44, 0.86]** | **<0.0001** |
| 6D – R1 | −0.156 | [−0.36, 0.05] | 0.226 |
| **6D – R2** | **0.278** | **[0.07, 0.49]** | **0.017** |
| 6D – R3 | −0.114 | [−0.32, 0.09] | 0.393 |
| C1 – C2 | 0.108 | [−0.10, 0.31] | 0.416 |
| **C1 – C3** | **0.940** | **[0.73, 1.15]** | **<0.0001** |
| C1 – R1 | 0.136 | [−0.08, 0.35] | 0.291 |
| **C1 – R2** | **0.569** | **[0.36, 0.78]** | **<0.0001** |
| C1 – R3 | 0.178 | [−0.04, 0.39] | 0.154 |
| **C2 – C3** | **0.832** | **[0.62, 1.05]** | **<0.0001** |
| C2 – R1 | 0.028 | [−0.19, 0.24] | 0.836 |
| **C2 – R2** | **0.462** | **[0.25, 0.67]** | **<0.0001** |
| C2 – R3 | 0.070 | [−0.14, 0.28] | 0.581 |
| **C3 – R1** | **−0.804** | **[−1.02, −0.59]** | **<0.0001** |
| **C3 – R2** | **−0.370** | **[−0.58, −0.16]** | **0.0018** |
| **C3 – R3** | **−0.762** | **[−0.98, −0.55]** | **<0.0001** |
| **R1 – R2** | **0.434** | **[0.22, 0.64]** | **0.0001** |
| R1 – R3 | 0.042 | [−0.16, 0.25] | 0.750 |
| **R2 – R3** | **−0.392** | **[−0.60, −0.18]** | **0.0005** |

*Table 2: Pairwise contrasts of estimated marginal means (log scale) from a negative binomial GLM of H. tvaerminnensis spore burden in Experiment 2.*
*P-values adjusted using the Benjamini–Hochberg method; 95% CI computed on log scale.*

| Contrast | Log difference (LRR) | 95% CI | adj. p |
|---|---|---|---|
| 1D – 2D | 0.154 | [−0.86, 1.17] | 0.787 |
| 1D – 6D | 0.461 | [−0.51, 1.43] | 0.466 |
| 1D – C1 | −0.410 | [−1.34, 0.52] | 0.466 |
| 1D – C2 | −0.441 | [−1.38, 0.50] | 0.466 |
| **1D – C3** | **5.229** | **[3.90, 6.56]** | **<0.0001** |
| 1D – R1 | 0.948 | [−0.116, 2.012] | 0.155 |
| **1D – R2** | **1.536** | **[0.350, 2.723]** | **0.037** |
| 1D – R3 | 0.646 | [−0.419, 1.711] | 0.366 |
| 2D – 6D | 0.308 | [−0.756, 1.372] | 0.642 |
| 2D – C1 | −0.563 | [−1.583, 0.457] | 0.403 |
| 2D – C2 | −0.594 | [−1.605, 0.417] | 0.390 |
| **2D – C3** | **5.076** | **[3.71, 6.44]** | **<0.0001** |
| 2D – R1 | 0.794 | [−0.363, 1.951] | 0.299 |
| 2D – R2 | 1.382 | [0.118, 2.646] | 0.083 |
| 2D – R3 | 0.493 | [−0.659, 1.645] | 0.466 |
| 6D – C1 | −0.871 | [−1.862, 0.120] | 0.155 |
| 6D – C2 | −0.902 | [−1.898, 0.094] | 0.155 |
| **6D – C3** | **4.768** | **[3.43, 6.11]** | **<0.0001** |
| 6D – R1 | 0.487 | [−0.631, 1.605] | 0.466 |
| 6D – R2 | 1.075 | [−0.145, 2.295] | 0.157 |
| 6D – R3 | 0.185 | [−0.933, 1.303] | 0.787 |
| C1 – C2 | −0.031 | [−0.978, 0.916] | 0.949 |
| **C1 – C3** | **5.639** | **[4.33, 6.95]** | **<0.0001** |
| **C1 – R1** | **1.358** | **[0.283, 2.433]** | **0.037** |
| **C1 – R2** | **1.945** | **[0.751, 3.139]** | **0.005** |
| C1 – R3 | 1.056 | [−0.020, 2.132] | 0.122 |
| **C2 – C3** | **5.670** | **[4.35, 6.99]** | **<0.0001** |
| **C2 – R1** | **1.389** | **[0.292, 2.486]** | **0.037** |
| **C2 – R2** | **1.976** | **[0.768, 3.184]** | **0.005** |
| C2 – R3 | 1.087 | [0.003, 2.171] | 0.120 |
| **C3 – R1** | **−4.281** | **[−5.70, −2.86]** | **<0.0001** |
| **C3 – R2** | **−3.694** | **[−5.13, −2.26]** | **<0.0001** |
| **C3 – R3** | **−4.583** | **[−6.02, −3.15]** | **<0.0001** |
| R1 – R2 | 0.588 | [−0.729, 1.905] | 0.466 |
| R1 – R3 | −0.302 | [−1.499, 0.895] | 0.678 |
| R2 – R3 | −0.889 | [−2.175, 0.397] | 0.299 |

*Table 3: Pairwise contrasts of estimated marginal means (log scale) from a negative binomial GLMM of O. colligata spore burden in Experiment 2.*
*P-values adjusted using the Benjamini–Hochberg method; 95% CI computed on log scale.*

| Contrast | Log difference (LRR) | 95% CI | adj. p |
|---|---|---|---|
| 1D – 2D | −0.711 | [−1.49, 0.07] | 0.122 |
| 1D – 6D | 0.817 | [−0.039, 1.67] | 0.106 |
| **1D – C1** | **−1.493** | **[−2.284, −0.702]** | **0.001** |
| 1D – C2 | −0.112 | [−0.936, 0.712] | 0.903 |
| **1D – C3** | **2.794** | **[1.466, 4.122]** | **0.0001** |
| 1D – R1 | −0.622 | [−1.432, 0.188] | 0.183 |
| 1D – R2 | −0.695 | [−1.533, 0.143] | 0.156 |
| 1D – R3 | −0.089 | [−0.890, 0.712] | 0.903 |
| **2D – 6D** | **1.528** | **[0.726, 2.331]** | **0.001** |
| 2D – C1 | −0.782 | [−1.510, −0.054] | 0.067 |
| 2D – C2 | 0.599 | [−0.168, 1.366] | 0.180 |
| **2D – C3** | **3.505** | **[2.184, 4.826]** | **<0.0001** |
| 2D – R1 | 0.089 | [−0.664, 0.842] | 0.903 |
| 2D – R2 | 0.016 | [−0.764, 0.796] | 0.969 |
| 2D – R3 | 0.622 | [−0.118, 1.362] | 0.156 |
| **6D – C1** | **−2.310** | **[−3.119, −1.501]** | **<0.0001** |
| 6D – C2 | −0.929 | [−1.846, −0.012] | 0.062 |
| **6D – C3** | **1.977** | **[0.648, 3.306]** | **0.009** |
| **6D – R1** | **−1.439** | **[−2.269, −0.609]** | **0.002** |
| **6D – R2** | **−1.513** | **[−2.370, −0.656]** | **0.002** |
| 6D – R3 | −0.906 | [−1.726, −0.086] | 0.062 |
| **C1 – C2** | **1.381** | **[0.608, 2.154]** | **0.001** |
| **C1 – C3** | **4.287** | **[2.999, 5.575]** | **<0.0001** |
| C1 – R1 | 0.871 | [0.111, 1.631] | 0.056 |
| C1 – R2 | 0.798 | [0.003, 1.593] | 0.086 |
| **C1 – R3** | **1.404** | **[0.656, 2.152]** | **0.001** |
| **C2 – C3** | **2.906** | **[1.599, 4.213]** | **0.0001** |
| C2 – R1 | −0.510 | [−1.308, 0.288] | 0.252 |
| C2 – R2 | −0.584 | [−1.417, 0.249] | 0.212 |
| C2 – R3 | 0.023 | [−0.764, 0.810] | 0.969 |
| **C3 – R1** | **−3.416** | **[−4.708, −2.124]** | **<0.0001** |
| **C3 – R2** | **−3.489** | **[−4.816, −2.162]** | **<0.0001** |
| **C3 – R3** | **−2.883** | **[−4.207, −1.559]** | **0.0001** |
| R1 – R2 | −0.074 | [−0.886, 0.738] | 0.909 |
| R1 – R3 | 0.533 | [−0.249, 1.315] | 0.219 |
| R2 – R3 | 0.607 | [−0.193, 1.407] | 0.183 |

*Figure 1: O. colligata infection rates in Experiment 2. Error bars represent 95% confidence intervals.*

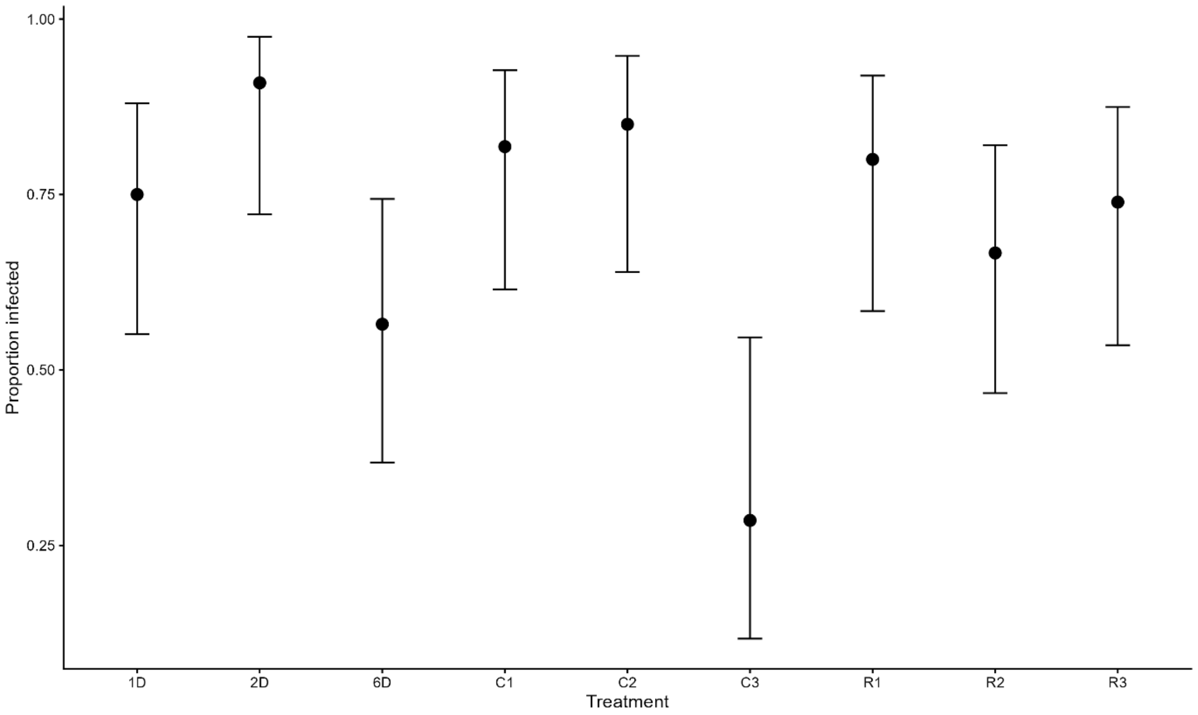


*Figure 2: H. tvareminnensis infection rates in Experiment 2. Error bars represent 95% confidence intervals.*

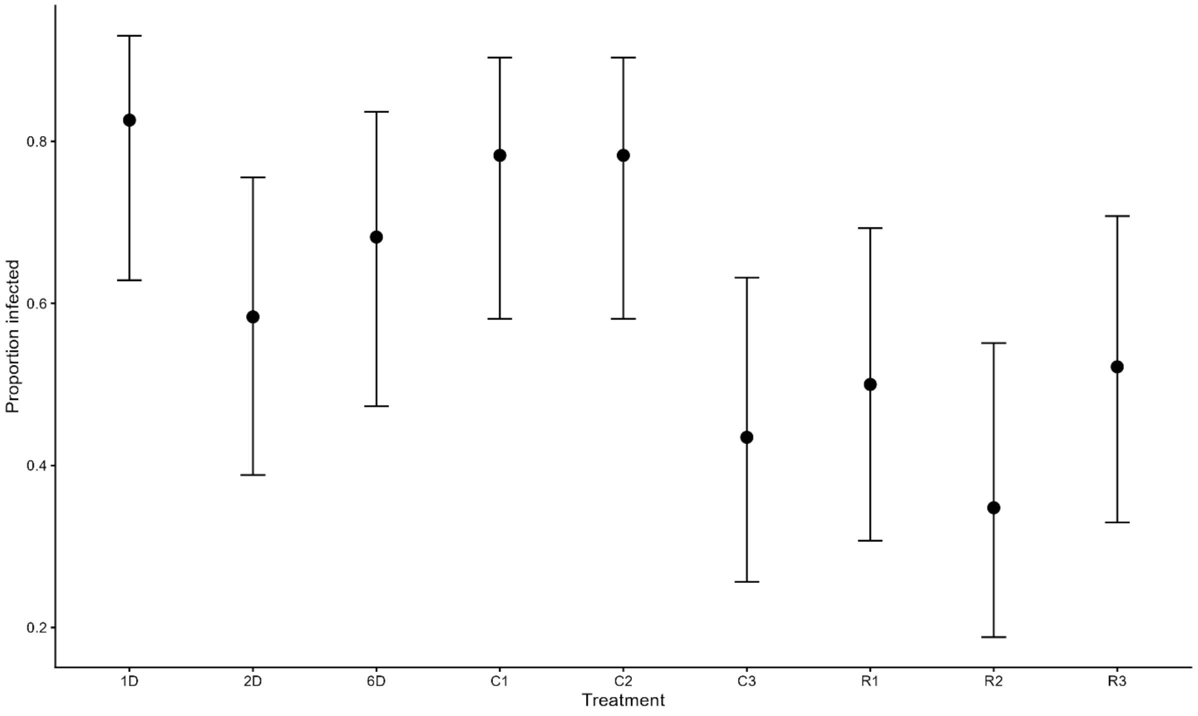
